# Building the Ferretome

**DOI:** 10.1101/014134

**Authors:** Dmitrii I. Sukhinin, Andreas K. Engel, Paul Manger, Claus C Hilgetag

## Abstract

Databases of structural connections of the mammalian brain, such as CoCoMac (cocomac.g-node.org) or BAMS (brancusi.usc.edu), are valuable resources for the analysis of brain connectivity and the modeling of brain dynamics in species such as the non-human primate or the rodent, and have also contributed to the computational modeling of the human brain. Another model species that is widely used in electrophysiological or developmental studies is the ferret; however, no systematic compilation of brain connectivity is currently available for this species. Thus, we have started developing a database of anatomical connections and architectonic features of the ferret brain (the Ferretome, www.ferretome.org). The main goals of this database project are: (1) to assemble structural information on the ferret brain that is currently widely distributed in the literature or in different in-house laboratory databases into a single resource which is open to the scientific community; (2) to try and build an extendable community resource that is beneficial to researchers in neuroinformatics and computational neuroscience, as well as to neuroanatomists, by adding value to their data through algorithms for efficient data representation, analysis and visualization; (3) to create techniques for the representation of quantitative and raw data; and (4) to expand existing database ontologies in order to accommodate further neuroarchitectural information for identifying essential relations between brain structure and connections.

The Ferretome database has adapted essential features of the CoCoMac methodology and legacy. In particular, its data model is derived from CoCoMac. It also uses a semantic parcellation of ferret brain regions as well as a logical brain maps transformation algorithm (objective relational transformation, ORT). The database is being developed in MySQL and has been populated with literature reports on tract tracing observations in the ferret brain using a custom-designed web interface that allows efficient and validated simultaneous input and proofreading by multiple curators. The database is also equipped with a web interface for generating output data that was designed with non-computer science specialist users in mind. This interface can be extended to produce connectivity matrices in several formats including a graphical representation superimposed on established ferret brain maps. An important feature of the Ferretome database is the possibility to trace back entries in connectivity matrices to the original studies archived in the system.

Currently, the Ferretome contains 50 reports on connections comprising 20 injection reports with more than 150 labeled source and target areas, the majority reflecting connectivity of subcortical nuclei. We hope that the Ferretome database will become a useful resource for neuroinformatics and neural modeling, and will support studies of the ferret brain as well as facilitate advances in comparative studies of mesoscopic brain connectivity.

## 1. Introduction

### Connectomics

A central perspective for imagining and analyzing brain data is the representation of neural relations as a complex network. This representation can be used for almost all structural-functional dimensions of the brain, from molecular to systems scales and structural to cognitive characterizations. The network graph representation is a powerful tool in the hands of neuroscientists, because it provides a formalized framework for the analysis of complex interactions (Klimm et al., 2014). Different types of brain connectivity can be distinguished, such as functional, effective and structural connectivity. In this context, functional connectivity indicates “temporal correlations between spatially remote neurophysiological events.” while effective connectivity reflects “the influence one neural system exerts over another” (Friston, 1994). The most fundamental type of connectivity is anatomical or structural connectivity, which provides a network basis of brain dynamics and function.

Several recent projects address the challenge of creating the complete structural network of the brain, the so-called connectome (Sporns et al., 2005), from the cellular to the mesoscopic and macroscopic scale. The neuronal micro-connectome, which is based on invasive methods of imaging and reconstruction of neuronal bodies (including synapses) from brain sections (see Van Essen et al., 2013 for an extended review), may be the ultimate structural basis of the brain. However, this approach faces many conceptual and technical challenges and so far has only been completed for the small nervous systems of the nematode *C. elegans*, possessing only 302 neurons (White et al., 1986; Varshney et al., 2011), as well as partly for zebrafish (Friedrich, 2013) and Drosophila (Chiang, 2011).

Examples for connectomes at the mesoscopic level include the recently published brain-wide mouse connectivity (Oh et al., 2014; Zingg et al., 2014), based on methods of optogenetics that label and trace axonal connections through areas of interests of the mouse brain. Further anatomical tracing techniques can be used to obtain structural connectivity at the mesoscopic and macroscopic level. The laborious method of histochemical tract-tracing has produced significant insights into the organization of brain connectivity and has resulted in an extensive body of connection data, for example, a detailed description and analysis of macaque monkey visual cortical connectivity (Felleman and Van Essen, 1991) and connectivity of the entire mesoscopic cat cortical (Scannel et al., 1995) and thalamocortical system (Scannel et al., 1999). These connectivity data were compiled from classical neuroanatomical studies that were assembled over many years. As a further attempt to systematize this approach of generating structural connectivity and in order to deal with methodological problems, such as different parcellation approaches and methods of labeling, connectivity databases such as the CoCoMac database were created (Stephan et al., 2001; Stephan, 2013; Bakker et al., 2012). Over a period of more than ten years, hundreds of tract-tracing reports for the macaque monkey brain have been collated in CoCoMac (Bakker et al., 2012).

A fundamental problem of conventional anatomical tract-tracing studies is that they cannot be performed in humans due to their invasiveness. This limitation raises questions about the applicability of data gathered in the animal models to humans. This problem can be ameliorated by comparative studies of different animal models (Bohland et al., 2009; Zingg et al., 2014), and through newly developed non-invasive techniques for imaging connectivity-related parameters. For example, diffusion imaging methods such as diffusion tensor imaging (DTI) or diffusion spectrum imaging (DSI) can be used to produce entire connectomes of a human brain in relatively short time (Van Essen et al., 2012). The method measures the anisotropy of water diffusion along axonal paths, which can then be used to infer the course of fiber tracts. The approach is systematically exploited by the Human Connectome Project (Toga et al., 2012), which aims to provide a comprehensive description of all long-range pathways of the human brain; however, the diffusion-based approaches are prone to several measuring and reconstruction artifacts (Farquharson et al., 2013).

The rise of new imaging methods such as DTI raises the question of whether we still need laborious and invasive anatomical track-tracing studies and associated connectivity databases. The answer is affirmative, as such conventional data provide a well-established 'ground truth' or 'gold standard' of structural brain connectivity. With this approach, one can directly observe the labeled origins and terminations of projection neurons in different brain regions, gather information on the axonal density and direction of projections as well as finer details such as the laminar origin and termination of projections. All of these aspects are currently not accessible with diffusion-based tractography. It should, however, be noted that the conventional anatomical tract-tracing studies are not without potential methodological problems either, considering, for example, mislabeling due to spillage of tracer injections into neighboring regions or the white matter (for more discussion of these issues see Kötter, 2001). Moreover, there are also problems associated with the many alternative ways of parcellating the brain into different areas, by not completely objectified criteria. In addition, many reports in the literature do not provide quantitative data about the number of labeled neurons or fine density of axonal terminations, but only binary information on the presence or absence of pathways, or comparative qualitative measures, such as ‘low’/ ‘average’/ ‘high’ density of connections (Lanciego and Wouterlood, 2011).

### The model system of the ferret brain

Due to limitations of directly investigating the structural connectivity of the human brain, research has turned to animals models, such as the ferret, where extensive developmental, behavioral or electrophysiological data can be obtained at low cost. One of the main advantages for the developmental studies is that the ferret brain is convoluted and the process of gyrification can be observed in detail (Sawada and Watanabe, 2012). Immaturity of the ferret at birth helps to observe processes that occur prenatally in other species, such as the cat, and conduct experiments with altered connectivity in order to observe the adaptation of cortical areas to new sensory stimuli (Noctor et al., 2001), how early lesions in one part of the brain may affect connectivity in other brain regions (Restrepo et al., 2002), and how early lesions may affect the development of topographical maps and connectivity between cerebral hemispheres (Restrepo et al., 2003). An additional advantage of the ferret model is that its brain shows substantial homologies with those of other species, such as the cat (Manger et al., 2010) as well as potentially with other carnivores such as the dog (Onishi et al., 2007). Taking these factors into account, extensive work has been performed in this model using electrophysiology to relate patterns of electrical activity with behavior (Bizley et al., 2013; Fritz et al., 2003). These mapping studies have shown that ferrets have highly complex sensory cortical systems (Innocenti et al., 2002; Manger et al., 2005; Phillips et al., 1988, Nelken and Versnel, 2000; Bizley et al., 2005; Bizley et al., 2007), making them an interesting model for the study of sensory processing pathways, response properties and topographies of sensory neurons and multisensory interactions. There exists no comparable model at the moment that combines elaborate and easily trainable behavior with the opportunity for extensive anatomical and physiological (as well as developmental) studies. In particular, similar studies in primates, which proceed in only very few labs, are much more restricted in the scope of investigations and the number of animals studied.

Hence, having a detailed macroconnectome of the ferret brain will facilitate comparative anatomical studies and support cross-domain exchange in anatomy, electrophysiology and connectomics; however, at the moment, no systematic compilation of connectivity is available for this species. Creating a repository of the macroconnectivity of the ferret brain is a complex task. The assembly of the data from published tract-tracing report faces similar problems as previously addressed by the CoCoMac database (Bakker et al., 2012) or projects such as BAMS (Bota et al., 2005) and neuroVIISAS (Schmitt and Eipert, 2012), see details below. In the following section, we provide a short review of existing database projects that aim at storing connectivity data, in order to define the parameters of a suitable architecture for the ferret brain connectivity database.

## 2. Comparable work

In the area of connectivity databasing, two main types of approaches for representing brain topography can be distinguished: coordinate-based versus semantic or logical parcellation schemes. The first type is represented by the XANAT system (Press et al., 2001), while the second approach is employed by the remainder of projects reviewed below.

XANAT (Press et al., 2001) was one of the first systems being developed for storing, comparing and analyzing the results of neuroanatomical connection studies. Data can be entered into the database by placing injection and label sites into canonical representations of the neuroanatomical structures of interest, along with verbal descriptions. After the entry procedure, a graphical search can be performed on the data by selecting a specific brain site or textual search with use of keywords or references to original studies. An important feature of the system is that data may be studied and compared relative to well-known neuroanatomical substrates or stereotaxic coordinates regardless of variable areal boundaries (Press et al., 2001). XANAT can be downloaded and run under Unix X window systems (reflected by the name of the system).

BAMS, the ‘Brain Architecture Management System’ (Bota et al., 2005), is a representative example of the attempt to store comprehensive structural descriptions of the brain. Information about four main entities and their attributes can be kept in the system: connections, relations, cell types and molecules. The connections entity represents records about data and metadata of macroscopic neuroanatomical projections between brain regions. The relations entity describes qualitative spatial relations between brain regions. Cell type attributes provide descriptions of neurons, neuronal population and their classifications. The molecules category represents data on molecules specific to neurons and brain regions.

BAMS is accessible online via a web interface (http://brancusi.usc.edu/bkms). The server part is written in PHP and the database itself is created in and handled by MySQL. In BAMS data can be stored and found for different species; however, the majority of it reflects structural descriptions of the rat. Some data can be exported for further analysis in structured formats (for example, as an adjacency matrix).

The NeuroVIISAS platform (NeuroVisualization, Image mapping, Information System for Analysis and Simulation) (Schmitt and Eipert, 2012) is an example of a neuroinformatics approach that attempts to link the storage of connectivity information with its visualization and analysis. NeuroVIISAS is an open framework in which one can perform integrative data analysis, visualization of the data and even population simulations (with the help of a link to popular software for neuronal simulations – NEST, see Gewaltig and Diesmann, 2007). During the data analysis step, it is possible to use many global network indices ranging from randomizations to the modified scale free network. Connectivity matrices can be visualized together with a display of indices, such as the clustering coefficient (Holland and Leinhardt, 1971) and joint degree distributions (Albert and Barabási, 2002). Visualization can be performed using the Paxinos and Watson atlas (Paxinos and Watson, 2006). Population simulations using the connectivity data can be performed using PyNEST (Davison et al., 2008) and NEST (Gewaltigand Diesmann, 2007). Neurobiologically defined connectivity is combined with computational neuroscience methods and after script generation and computations, finished simulation results can be imported back into NeuroVIISAS and visualized again in various ways, including 3D visualization. NeuroVIISAS is free software implemented in Java with versions for Windows and Linux, which can be operated locally. The main advantage of this approach is that a researcher's own data (connectivity or mapping) can be quickly added to the framework and analyzed, visualized and simulated locally.

Finally, CoCoMac (Collation of Connectivity data on the Macaque brain) is a connectivity database and neuroinformatics platform that has been developed for more than a decade (Stephan et al., 2001; Bakker et al 2012; Stephan, 2013). CoCoMac aims to store two main modalities of data: connectivity - tract tracing studies as well as mapping studies of (mainly) rhesus macaque. CoCoMaC faces many challenges, such as the absence of spatial coordinates and of a universally accepted brain map for the Macaque, inconsistency of alternative brain parcellation schemes, as well as ambiguity and contradictions of results from different tract-tracing studies. The CoCoMac creators postulated five main principles for their project: Objectivity, Reproducibility, Transparency, Flexibility and Simplicity. These principles reflect the way in which to treat links of original data, to insert data as well as the processing of this data. To meet the demands of these principles, a special algorithmic framework was developed, called Objective Relational Transformation (ORT; Stephan and Koetter, 1999; Stephan et al., 2000). In short, this framework allows the transformation of all available connectivity data of one brain map into another according to relations between areas and brain maps established in different papers using an encoding of logico-spatial relations between them (i.e. some area A is smaller than/bigger than/equal with/overlaps with some area B).

Originally, CoCoMac was created in MS Access, but subsequently the database was converted to MySQL and made accessible through a web interface, with the server side programmed in PHP. With the recent update to new version (at a new address, http://cocomac.g-node.org) CoCoMac received several new features including a search/browse wizard and direct access to the content of database via specifically developed viewers (Bakker et al., 2012).

In this section we touched upon the modern approach in neuroscience to the issue of representing brain connectivity with use of graph models at different scales. Despite the rise of new methods such as DSI/DTI, at the macroscopic level, anatomical tract-tracing studies are still the most reliable source of data for graphic representation of the brain. Presence of macroscopic connectivity data for as many species as possible will facilitate comparative studies and deepen our understanding of the human brain. One popular animal model is the ferret due to its valuable features such as gyrification and immaturity at birth. Creating a complete brain connectivity scheme of an animal even as small as a ferret is a complex task that requires the help of modern methods in computer science such as online databasing. In the next section of this paper we turn to the issue of building such database, populating it with data, supporting it and extracting results.

## 3. Methods

From a schematic point of view, the main structure of the database was derived from the CoCoMac project (Stephan et al., 2001). The CoCoMac data model allows the storage of data relating to main entities such as literature references, brain map descriptions, injection of tracers and labeled sites. One of the key concepts of CoCoMac is an approach for specifying the reliability of the data by Precision Data Codes (PDC). PDCs were used in CoCoMac in order to cope with situations when the text of a paper contradicts figures or/and tables. PDC is coded by different letters from “A” to “Q”, where “A” is the most reliable and consistent description. For example, the PDC code “A” for specifying a brain area means that “The area is named explicitly in the text/tables and identified with certainly. Additional figures explicitly support the text by showing present (or missing) label in areas defined by names and/or borders”, where “Q” means: “The information about the (un)labeled area is not from an original research report, but from a review article” (more details in Stephan et al., 2001). CoCoMac provides several types of PDC's for different types of data, for example, PDC_BrainArea, PDC_lamina with their own descriptions.

This database (DB) schema was taken with adjustments made according to ferret species-specificity and additional requirements established during the survey and conceptual planning. For example, CoCoMac has the means to store data about subspecies of Macaque, which was unnecessary here. The main difference, however, was an introduction of extensible and flexible tables that store data about ferret brain architecture. For this novel type of data, the same PDC method of specifying the data reliability was employed. Different aspects of PDC_Architecture were gathered from the literature and can be used for an entire brain area as well as for area subcompartments, such as individual cortical layers. For example, DB collators can specify types of neurons in a layer and provide PDC for this entry.

Another distinct feature that was adopted from CoCoMac is the approach of Objective Relational Transformation (ORT). This powerful routine allows the automatic conversion of all available data (including PDCs) from one given brain map to another. ORT uses a custom-developed relational algebra that describes five main relations between areas: identical, subarea, larger, overlap, disjoint (for details see Stephan et al., 2000). Specifically, this means that if there exists a report that specifies a relation among brain maps, then it is possible to transform connectivity data from one report to another and hence to build a comprehensive description of ferret brain connectivity. For example, if two areas from two different brain maps are specified by a report as “identical”, then all data associated with these areas can be transferred from one map to another. In addition to transforming data for known relations among brain maps, ORT is capable of discovering previously undefined relations between brain areas of different brain maps (i.e., not specified in any tract tracing report). For example, if it is known that “A” is identical to “B” and “B” identical to “C”, it can be inferred that “A” is identical to “C”. The algorithm can also identify inconsistent relations (such as that “A” is bigger than “B” while also “B” is bigger than “A”). In Ferretome.org, the implementation of ORT serves not only the same function as in CoCoMac for converting connectivity information among brain maps, but it is used as well for transferring architecture data from one area to another.

Going deeper into technical details, ferretome.org represents a typical web application with a front-office and a back-office that is supported by a database. As a database management system, the reliable and free MySQL (www.mysql.com) was employed and phpMyAdmin (www.phpmyadmin.net) was used to handle the initial creation and editing of tables.

According to the data model, the input interface (back-office) was developed using PHP for server side operations and the bind of html5 and JavaScript for the user side. To simplify development, some parts of open source CodeIgniter PHP framework (https://ellislab.com/codeigniter) were used, in particular DB manipulation libraries. The interface guides users step-by-step through the process of the data input and data validation and supplies them with hints about types of data and extensive help with searching for related or previously inserted data by means of autocompletion.

To successfully complete the project, a conveyor was created with four main steps: (1) discovery of tract tracing reports, (2) short-listing them and putting them into queue for input, (3) input by one DB collator, (4) proofreading by another DB collator. To allow the work of several DB collators simultaneously, the conveyor-like system was integrated into the DB interface and data model, by means of an input wizard. This allows users to select new tasks and keep track of their current tasks.

Fig. 2. Tasks selection and management menu as well as short progress report.

A major difference between the ferret connectivity DB and CoCoMac is that a special interface for the input of architectural data was developed. For the convenience of the users, this interface mostly utilizes the JavaScript ability to manipulate the contents of the web page, which allowed the user, in a couple of clicks, to assign all necessary architectural features to a brain area or to a subregion (layer/part) of a brain area.

Although the input interface allows navigation across already inserted data, for the convenience of the end users, an entire new interface for data browsing was created (front-office). It uses the same technologies and interacts with the database in read-only mode. The data browsing interface provides different means of searching information and creating summaries of stored data.

For the purpose of integration with connectivity analysis software and neuronal simulator packages, an interface for connectivity output was created. This interface generates adjacency matrices that can be saved in different formats. Ferretome.org automatically maps all available data into a selected map using the interpretation of the ORT algorithm.

### Results

Currently, Ferretome.org can be characterized as a beta version. While it integrates all connectivity information for the ferret currently available in the literature, the available information itself is sparse, so the information contained in the Ferretome about the brain architecture and macroconnectome of the ferret brain is still limited. This limitation arises from the relatively small number of anatomical connectivity reports published so far on the ferret, the larger part of which covers subcortical connections; however, the database is continuously being populated with newly appearing reports. Already stored records can be accessed via the web interface, where the full summary of inserted data for a given publication is represented as a table. This table can be dynamically extended to display links with other publications (e.g., if brain maps were defined in a different paper and the current record is using their parcellation scheme to provide tract tracing results).

Using the same interface, the architecture of the brain areas can be obtained directly from the extracted data of a paper, as well as from another records by using an ORT algorithm that transforms connectivity data from one area to another, if relations among parcellations schemes were specified.

At the current point, more than 150 ferret brain papers were reviewed, 50 them were collated into the database and for 30 of them, that contain mapping or connectivity data, the proofreading is finished. These 30 reports contain 20 unique injections sites with 200 labelling sites in both ipsi- and contra-lateral hemispheres of the ferret brain. Architectural data is currently provided for 12 distinct brain areas, primarily for visual and auditory cortex.

## 4. Discussion and outlook

Differences in the techniques of different neuroanatomical labs, and the absence of well-established standards for producing tract-tracing reports, create challenges in extracting data for systematic computational analysis and cross-species comparative studies. After a review of existing technologies, approaches and methods, it appeared that the most suitable approach for databasing structural information of the ferret would be a CoCoMac-like approach and schema. The reasons for this decision were similar to those of the initial CoCoMac development. First, most tract-tracing reports do not provide the exact spatial location of injections sites but rather semantic positions (such as an injection being made into ‘primary visual cortex’ or ‘area 17’). Second, brain areas in one brain map can have different boundaries in another brain map. In order to build a comprehensive description of ferret brain connectivity, one needs means to relate one brain area and its connectivity in one parcellation to another brain area in a different parcellation. This transformation is tedious and error-prone to perform by hand, and hence requires automation. Thus we focused on the problem of how to apply and how to improve the CoCoMac approach to the case of the ferret. Our system includes the main features of the CoCoMac approach, including PDC coding and the ORT algorithm, but in addition, we have extended the database schema in order to flexibly accommodate the representation of architecture of brain areas.

To provide a wide base for the subsequent use of the database, several additional structural parameters were included. The first of them was taken from the notion that connectivity appears to be closely related to the architectonic similarity of potentially connected areas (e.g., Hilgetag and Grant, 2010; Beul et al., 2014). Many tract-tracing reports also provide descriptions of brain cytoarchitecture. This description includes the classification of cells, number of layers and sublayers and their density amongst other features. Such cytoarchitectonic descriptions encounter the same problem as connectivity data, because they are usually defined by researchers within their own brain map and hence need to undergo the transformation from one brain map to another.

An important extension of the CoCoMac methodology is to link connectivity data to analysis and simulation tools. This approach is vital not only for understanding functional implications of connectivity, but also for validating data inserted into the database by providing summarized feedback that can be compared to global models of connectivity organization. Hence the connectivity database should be easily interfaced to analysis tools (for example, Brain Connectivity Toolbox, www.brain-connectivity-toolbox.net) and simulation packages (e.g., The Virtual Brain, www.thevirtualbrain.org), and should provide a multitude of popular input and output formats (XML, JSON, etc.) for the use of external packages. As a consequence, the Ferretome is set up to provide such interfaces.

On the practical side, an efficient implementation and management system is required in order to maintain an up-to-date connectivity database that is quick and functional as well as easy to handle by administrators and users. One way of achieving this aspect is by providing constant web access to all parts of database. In this case data in the database can be reviewed not only by the database collators, but also external experts.

Although in the current state our database does not contain sufficient data to provide connectivity and architectural data for the entire ferret brain, it may be sufficient to identify underrepresented brain areas where, for various reasons, tract-tracing studies have not yet been conducted. As soon as new tract-tracing reports will appear in literature, the data will be added to ferretome.org.

The collated data do not have to be restricted to cortical connectivity and point-to-point connection systems, but could also include the connectivity of neuromodulatory systems. These systems typically have localized cell populations (such as the orexinergic neurons in the hypothalamus, or the cholinergic and noradrenergic neurons in the pons) that project widely throughout the brain and spinal cord (Dell et al., 2013). These projections are easily identified with immunohistochemistry, and could be readily plotted and quantified with stereological techniques (in terms of regional densities, distribution by cortical layers and neuronal types, etc.) and added to the database. In addition, the distribution, in a quantified manner, could be determined for the GABAergic neurons stained with parvalbumin, calbindin and calretinin. This effort would provide a unique insight into the inhibitory systems within the brain, which are often ignored in favor of the excitatory ones.

In addition to storing fundamental connectivity and architectural data for the ferret brain, several additions are planned for ferretome.org that will make access to the data easier or more functional. In the short term, we are planning the integration of visualization tools that can be deployed at the users' computer clients (directly in a browser); for example, with the use of WebGL technology which will allow future integration with a planned atlas of the ferret brain. Taking into account that Ferretome has an additional data modality that represents the architecture of brain areas, the idea of visualization tools should be extended. This visualization tool should give to users the possibility to observe simultaneously connectivity data and architectural data (see fig. 5). By analogy with connectivity data, researchers should have the ability to perform a quick analysis of architectural data right in their browser. Definitely, it will be helpful if architectural information, for example, on the cellular density and thickness of cortical layers, can be read out in standard formats for further offline analysis.

**Fig. 1.**
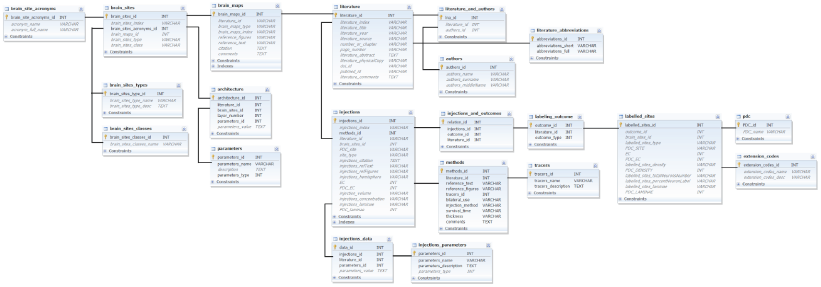
Simplified schema of the Ferretome connectivity database. Only main links and tables are shown.

**Fig. 2.**
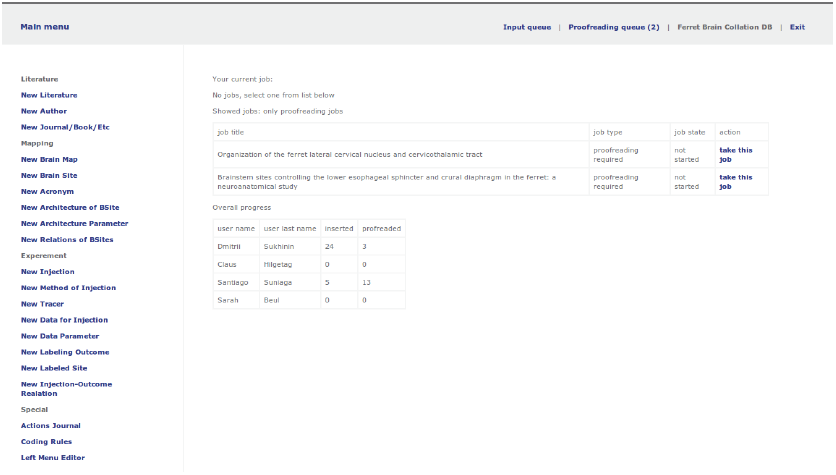
Tasks selection and management menu as well as short progress report.

**Fig. 3.**
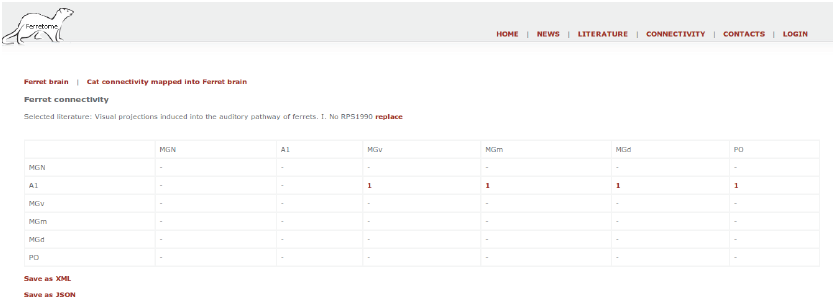
The connectivity output interface of the ferretome.org

**Fig. 4.**
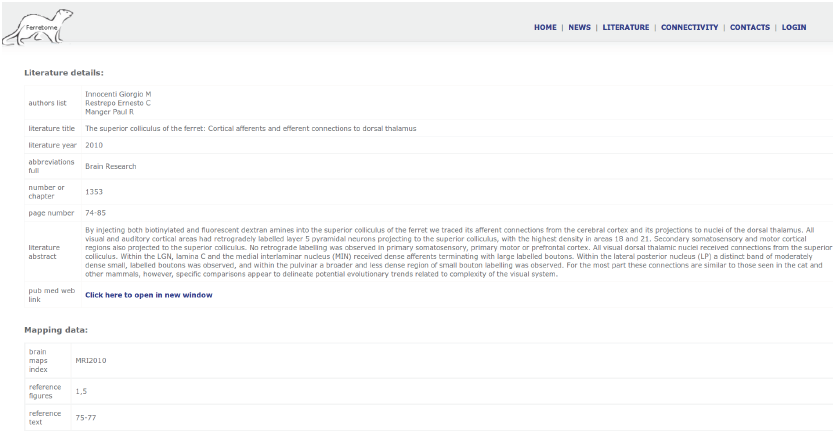
Summary for inserted data of one paper

**Fig. 5.**
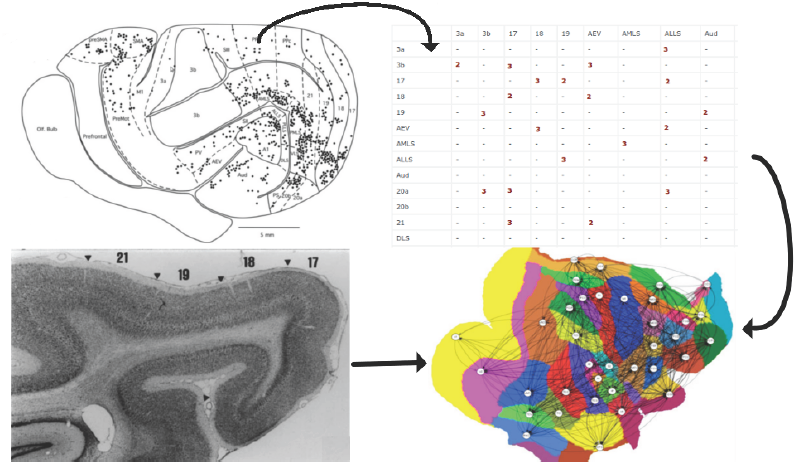
Data integration and visualization concept. Through using a web interface, both connectivity and architectural information can be obtained from the database and jointly visualized in a brain map. Credit: left side - (Manger et al., 2010), Upper right - ferretome.org (not actual data), bottom right (Bakker et al., 2010).

Moreover, following the example of the NeuroVIISAS platform (Schmitt and Eipert, 2012), integration with connectivity analysis tools, such as the BrainConnectivityToolbox (www.brain-connectivity-toolbox.net), or tools for modeling brain dynamics, like TheVirtualBrain (www.thevirtualbrain.org; Ritter et al., 2013), will be provided. This integration will allow to characterize features of structural nodes and circuits and link them to aspects of brain dynamics and function.

In the long-term, an important goal is the involvement of the scientific community, in particular of experimental neuroanatomists, for contributing new data or validating the data already existing in the database. This step is essential for verifying the overall consistency of the data and facilitating the dialog among all parties who are interested in ferret brain structure and function. Thus, the system has to be designed in such a way that it is accessible and appealing to experimentalists studying the ferret brain. Based on this idea of community participation, one of the potentials to increase the value of this databasing project is to have the ability to store the raw data (images, quantitative information) taken directly from experiments. From the technological point of view, this is a challenging task that requires development of special storage subsystems and algorithms for data access as well as data protection methods at different levels of data access, public and private. As a further extension of the concept of this connectivity database, we also consider the possibility of adding the modality of large-scale functional connectivity of the ferret brain, both at rest and during a task. This idea can be implemented with the same methodology as CoCoMac and Ferretome.org, by providing information on the reliability of data and by transformation of data across different brain maps. Ultimately, the structural and functional perspective of such data can be linked through computational modeling platforms.

Certainly, ferretome.org should have the functional capacity to extract data of all modalities (including computed brain maps relations) in different formats in order to be integrated with different software such as The Virtual Brain and including various (online) atlases. Integration with atlases will be useful not only as a new perspective, but can provide new knowledge in the area of comparative studies. For example, co-registering connectivity data with the SVG based Common Atlas format developed by (Majka et al., 2012) has helped to facilitate studies in a variety of species.

In summary, here we introduced Ferretome.org, a ferret brain macro-connectivity and architecture database. This project is built upon the experience of a previous generation of neuroinformatics project such as XNAT, BAMS, NeuroVIISAS and in particular CoCoMac. In particular, ferretome.org inherited from CoCoMac the basic methodology and philosophy of objectivity and reproducibility, and follows the same data collation rules and standard. In addition, we extended basic CoCoMaC methodology in order to capture architectural data that provides an important context for the connectivity data. Currently, we are moving towards extensive population of the database with newly published results and hope to provide a useful contribution to studying ferret brain structure and function.

## Acknowledgements

We gratefully acknowledge financial support by the DFG, SFB 936, projects A1 (authors DIS and CCH) and A2 (AKE).

## Literature

Albert, R., & Barabási, A. L. (2002). Statistical mechanics of complex networks. Reviews of Modern Physics, 74(1), 47.

Chiang, S. (2011). Imaging atoms and molecules on surfaces by scanning tunnelling microscopy. Journal of Physics D: Applied Physics. Doi:10.1088/0022-3727/44/46/464001

Bakker, R., Larson, S. D., Strobelt, S., Hess, A., Wójcik, D. K., Majka, P., & Kötter, R. (2010). Scalable brain atlas: from stereotaxic coordinate to delineated brain region. Frontiers in Neuroscience, 4(5).

Bakker, R., Wachtler, T., & Diesmann, M. (2012). CoCoMac 2.0 and the future of tract-tracing databases. Frontiers in NeuroinformaticsNeuroinformatics, 6.

Beul, S. F., Grant, S., & Hilgetag, C. C. (2014). A predictive model of the cat cortical connectome based on cytoarchitecture and distance. Brain Structure and Function, 1–18, 10.1007/s00429-014-0849-y.

Bizley, J.K., Nodal, F.R., Nelken, I., King, A.J. (2005) Functional organization of ferret auditory cortex. Cereb Cortex 15: 1637–1653.

Bizley, J.K., Nodal, F.R., Bajo, V.M., Nelken, I., King, A.J. (2007) Physiological and anatomical evidence for multisensory interactions in auditory cortex. Cereb Cortex 17: 2172–2189.

Bizley, J.K., Walker, K.M., Nodal, F.R., King, A.J., Schnupp, J.W. (2013) Auditory cortex represents both pitch judgments and the corresponding acoustic cues. Curr Biol. 23(7):620–625. doi: 10.1016/j.cub.2013.03.003.

Bohland, J. W., Wu, C., Barbas, H., Bokil, H., Bota, M., Breiter, H. C., ... & Mitra, P. P. (2009). A proposal for a coordinated effort for the determination of brainwide neuroanatomical connectivity in model organisms at a mesoscopic scale. PLoS Computational Biology, 5(3), e1000334.

Bota, M., Dong, H. W., & Swanson, L. W. (2005). Brain architecture management system. Neuroinformatics, 3(1), 15–47.

Davison, A.P., Brüderle, D., Eppler, J.M., Kremkow, J., Muller E., Pecevski, D.A., Perrinet L. and Yger P. (2008) PyNN: a common interface for neuronal network simulators. Front. Neuroinform. 2:11 doi:10.3389/neuro.11.011.2008

Dell, L.A., Kruger, J.L., Pettigrew, J.D., Manger, P.R. (2013) Cellular location and major terminal networks of the orexinergic system in the brain of two megachiropterans. J Chem Neuroanat. 53:64–71. doi: 10.1016/j.jchemneu.2013.09.001.

Farquharson, S., Tournier, J. D., Calamante, F., Fabinyi, G., Schneider-Kolsky, M., Jackson, G. D., & Connelly, A. (2013). White matter fiber tractography: why we need to move beyond DTI: Clinical article. Journal of Neurosurgery, 118(6), 1367–1377.

Fritz, J., Shamma, S., Elhilali, M., Klein, D. (2003) Rapid task-related plasticity of spectrotemporal receptive fields in primary auditory cortex. Nat Neurosci. 6(11): 1216–1223.

Felleman, D. J., and Van Essen, D. C. (1991). Hierarchical processing in the primate cerebral cortex. Cereb. Cortex 1, 1–47. doi: 10.1093/cercor/1.1.1

Friedrich, R. W. (2013). Neuronal computations in the olfactory system of zebrafish. Annual Review of Neuroscience, 36, 383–402. doi:10.1146/annurev-neuro-062111-150504

Friston, K. J. (1994). Functional and effective connectivity in neuroimaging: a synthesis. Human Brain Mapping, 2, 56–78

Gewaltig, M.O. & Diesmann, M. (2007). NEST (Neural Simulation Tool) Scholarpedia 2(4):1430.

Hilgetag, C.C., Grant, S. (2010). Cytoarchitectural differences are a key determinant of laminar projection origins in the visual cortex. NeuroImage 51:1006–1017. doi: 10.1016/j.neuroimage.2010.03.006

Holland, P. W., & Leinhardt, S. (1971). Transitivity in structural models of small groups. Comparative Group Studies.

Innocenti. G.M., Manger, P.R., Masiello, I., Colin, I., Tettoni, L. (2002) Architecture and callosal connections of visual areas 17, 18, 19 and 21 in the ferret (Mustela putorius). Cereb Cortex 12: 411–422

Kötter, R. (2001). Neuroscience databases: tools for exploring brain structure-function relationships. Philosophical Transactions of the Royal Society of London. Series B, Biological Sciences, 356, 1111–1120. doi:10.1098/rstb.2001.0902

Klimm, F., Bassett, D. S., Carlson, J. M., & Mucha, P. J. (2014). Resolving structural variability in network models and the brain. PLoS Computational Biology, 10(3), e1003491. doi:10.1371/journal.pcbi.1003491

Lanciego, J. L., & Wouterlood, F. G. (2011). A half century of experimental neuroanatomical tracing. Journal of Chemical Neuroanatomy, 42(3), 157–183.

Majka, P., Kublik, E., Furga, G., & Wójcik, D. K. (2012). Common atlas format and 3d brain atlas reconstructor: infrastructure for constructing 3d brain atlases. Neuroinformatics, 10(2), 181–197.

Manger, P.R., Engler, G., Moll, C.K.E., Engel, A.K. (2005) The anterior ectosylvian visual area of the ferret: a homologue for an enigmatic visual cortical area of the cat? Eur J Neurosci 22: 706–714

Manger, P. R., Restrepo, C. E., & Innocenti, G. M. (2010). The superior colliculus of the ferret: Cortical afferents and efferent connections to dorsal thalamus. Brain Research, 1353, 74–85. doi:10.1016/j.brainres.2010.07.085

Nelken, I., Versnel, H. (2000) Responses to linear and logarithmic frequency-modulated sweeps in ferret primary auditory cortex. Eur J Neurosci 12: 549–562

Noctor, S. C., Palmer, S. L., McLaughlin, D. F., & Juliano, S. L. (2001). Disruption of layers 3 and 4 during development results in altered thalamocortical projections in ferret somatosensory cortex. Journal of 21(9), 3184–95

Oh, S. W., Harris, J. a, Ng, L., Winslow, B., Cain, N., Mihalas, S., ... Zeng, H. (2014). A mesoscale connectome of the mouse brain. Nature, 508, 207–14. doi:10.1038/nature13186

Onishi, T., Mori, T., Yanagihara, M., Furukawa, N., & Fukuda, H. (2007). Similarities of the neuronal circuit for the induction of fictive vomiting between ferrets and dogs. Autonomic Neuroscience, 136(1), 20–30.

Paxinos, G., & Watson, C. (2006). The rat brain in stereotaxic coordinates: hard cover edition. Academic Press.

Phillips, D.P., Judge, P.W., Kelly, J.B. (1988) Primary auditory cortex in the ferret (Mustela putorius): neural response properties and topographic organization. Brain Res 443: 281–294

Press, W. A., Olshausen, B. A., & Van Essen, D. C. (2001). A graphical anatomical database of neural connectivity. Philosophical Transactions of the Royal Society of London. Series B: Biological Sciences, 356 (1412), 1147–1157.

Ritter, P., Schirner, M., McIntosh, A. R., & Jirsa, V. K. (2013). The virtual brain integrates computational modeling and multimodal neuroimaging. Brain Connectivity, 3(2), 121–145.

Restrepo, E., Manger, P., Innocenti, G. (2002) Retinofugal projections following early lesions of the visual cortex in the ferret. Eur. J. Neurosci. 16: 1713–1719.

Restrepo, C., Manger, P., Spenger, C., Innocenti, G. (2003) Immature cortex lesions alter retinotopic maps and interhemispheric connections. Ann. Neurol. 54: 51–65

Sawada, K., & Watanabe, M. (2012). Development of cerebral sulci and gyri in ferrets (Mustela putorius). Congenital Anomalies, 52(3), 168–175.

Scannell, J. W., Blakemore, C., & Young, M. P. (1995). Analysis of connectivity in the cat cerebral cortex. Journal of Neuroscience, 15, 1463–1483.

Scannell, J. W., Burns, G. A. P. C., Hilgetag, C. C., O’Neil, M. A., & Young, M. P. (1999). The connectional organization of the cortico-thalamic system of the cat. Cerebral Cortex, 9, 277–299. doi:10.1093/cercor/9.3.277

Schmitt, O., and Eipert, P,. (2012). NeuroVIISAS: approaching multiscale simulation of the rat connectome. Neuroinformatics 10.3 : 243–267.

Sporns O, Tononi G, Kötter R (2005) The Human Connectome: A Structural Description of the Human Brain. PLoS Comput Biol 1(4): e42. doi:10.1371/journal.pcbi.0010042

Stephan K. E., Kötter R (1999). One cortex - many maps: An introduction to coordinate-independent mapping by Objective Relational Transformation (ORT). Neurocomputing 26-27: 1049–1054

Stephan, K. E., Zilles, K., & Kötter, R. (2000). Coordinate–independent mapping of structural and functional data by objective relational transformation (ORT). Philosophical Transactions of the Royal Society of London. Series B: Biological Sciences, 355(1393), 37–54.

Stephan, K. E., Kamper, L., Bozkurt, A., Burns, G. A., Young, M. P., & Kötter, R. (2001). Advanced database methodology for the Collation of Connectivity data on the Macaque brain (CoCoMac). Philosophical Transactions of the Royal Society of London. Series B: Biological Sciences, 356(1412), 1159–1186.

Stephan, K.E. (2013). The history of CoCoMac. NeuroImage 80: 46–52.

Toga, A. W., Clark, K. A., Thompson, P. M., Shattuck, D. W., & Van Horn, J. D. (2012). Mapping the human connectome. Neurosurgery, 71(1), 1.

Van Essen, D. C., Ugurbil, K., Auerbach, E., Barch, D., Behrens, T. E. J., Bucholz, R., ... & Yacoub, E. (2012). The Human Connectome Project: a data acquisition perspective. NeuroImage, 62(4), 2222–2231.

Van Essen, D. C., Smith, S. M., Barch, D. M., Behrens, T. E., Yacoub, E., & Ugurbil, K. (2013). The WU-Minn human connectome project: an overview. NeuroImage, 80, 62–79.

Varshney, L., Chen, B., Paniagua, E., Hall, D., & Chklovskii, D. (2011). Structural properties of the Caenorhabditis elegans neuronal network. PLoS Computational Biology, 7, e1001066. doi:10.1371/journal.pcbi.1001066

White J. G., Southgate E., Thomson J.N., and Brenner S. The structure of the nervous system of the nematode caenorhabditis elegans. Phil. Trans. Royal Soc. London. Series B, Biol Scien. 314, 1–340.

Zingg, B., Hintiryan, H., Gou, L., Song, M. Y., Bay, M., Bienkowski, M. S., ... Dong, H. W. (2014). Neural networks of the mouse neocortex. Cell, 156, 1096–1111. doi:10.1016/j.cell.2014.02.023.

